# Validation of the in-house designed one-trial inhibitory avoidance test for two strains of adult zebrafish

**DOI:** 10.1101/2023.05.05.539645

**Authors:** Jorge M. Ferreira, Joana Silva, Sofia Barros, Inês Caetano, Pedro Fernandes, Anna Olsson, Ana M. Valentim

**Affiliations:** i3S - Instituto de Investigação e Inovação em Saúde, Universidade do Porto, Portugal; IBMC - Instituto de Biologia Molecular e Celular (IBMC), Universidade do Porto, Portugal; Instituto de Ciências Biomédicas Abel Salazar (ICBAS-UP), Universidade do Porto; Faculdade de Ciências da Universidade do Porto (FCUP), Universidade do Porto; Faculdade de Engenharia da Universidade do Porto (FEUP), Universidade do Porto

**Keywords:** Zebrafish, memory, validation, behaviour, open-source

## Abstract

Open-source validated tools are important for affordable high-quality science. Zebrafish model is often used in behavioral sciences, thus validated tools must be developed for this species. We aimed to implement the memory task one-trial inhibitory avoidance test in our laboratory by creating a custom-built 3D printed apparatus and custom-made hardware and software using microcontrollers; this decreases the costs and increases methodological flexibility. In this task, a mild electric shock (3.3 ± 0.3V and 2A for 5sec) is used as an aversive stimulus for the one-trial inhibitory avoidance task based on classical conditioning. For this study we used 72-adult zebrafish from the AB and TU strain. The aversive stimulus caused a robust and long-lasting memory that was learned in one session. The apparatus consisted in one aquarium with two compartments (white and black) where, during the conditioning session, the shock was delivered in the preferred side (black), followed by 30min of immersion in 20μM MK-801, a compound known for inducing amnesia (amnesic group, n= 20 for AB and 17 for TU), or in clean water (control, n= 20 for AB and 15 for TU). Both the AB and TU strain animals from the control group took more time to enter in the black compartment after conditioning (p≤0.039), indicating avoidance for the black side, while the latency was not altered in the MK-801 group, demonstrating impaired memory. In summary, this open-source apparatus is an affordable and validated option for memory assessment for the two most used strains of zebrafish in research.

## 1. Introduction

Creating, refining, and replicating protocols are key components of science. With the increase of zebrafish (*Danio rerio*) use as an animal model, accounting for 5% of all the animals used in the EU [1], there is an additional need for new and validated protocols that can be replicated. For a method to be replicable, it is crucial that it is properly reported in the literature, with all the details described. Failing to do so leads to the reproducibility crisis [2] and the waste of animals, which is an issue for the scientific results and for the 3Rs [3].

Often behavioral protocols tested in rodents are adapted to zebrafish, as they have become a relevant model in different areas of research. It is rather important to correctly adapt from rodents to zebrafish as they are different species with totally different needs. Such example is when adapting the white/black protocol from rodents to zebrafish, where the use of incorrect terminology can lead researchers to infer preferences incorrectly [4]. One example of this is the memory task passive avoidance [5]. In this test, also known as inhibitory avoidance test [6], the animal learns to avoid an unpleasant stimulus by limiting its locomotion and exploration.

For zebrafish, this test was adapted [7] using a black and white box where the animal will receive a mild shock in the black side of the box (their preferred side [8]) which they should learn to later avoid. To successfully replicate this task, a complete report of the shock profile is needed. The problem is that, although several articles report the voltage used and the duration of the shock, no report on the intensity of the shock (current in Amperes) was present. This is problematic as a shock relies on four major components: intensity, current, duration and type of current [9]. If information on one of these is missing, it is impossible to assess how strong the shock was and impossible to replicate the method. Also, for the components that are reported in the literature there is great variation, with voltages ranging from 3 and 12 volts, times under shock from 1ms to 5000ms and currents being AC or DC [7, 10-15], thus standardization is required. Another issue for implementing this task is the lack of a commercial version of the apparatus; and the ones that could be adapted were very expensive. Therefore, our objective was to create an affordable open source custom-made apparatus that allowed testing the memory of the two most used strains of zebrafish (AB and TU [16]), validating the apparatus and the protocol. Indeed, different strains have been describe to exhibit different behaviors [15, 16].

The validation of the test was pharmacological, using as a known amnesic as a positive control, the NMDA receptor antagonist MK-801. As an amnesic drug [17, 18], animals treated with this substance are expected to show memory impairment, indicating that the test measures what it is supposed to evaluate, memory.

## 2. Material and methods

### 2.1. Ethics statement

All procedures were carried out under animal facility, personal and project licenses approved by the National Competent Authority for animal research (Direção-Geral de Alimentação e Veterinária, Lisbon, Portugal) (approval number: 014703), and by the Animal Welfare and Ethics Review Body of the Institute for Research and Innovation in Health, according to the European Directive 2010/63/EU on the protection of animals used for scientific purposes, and its transposition to the Portuguese law, ‘Decreto Lei’ 113/2013.

### 2.2. Animals and Housing

Sample size calculation was performed in G*Power 3.1 (University of DulJsseldorf, Germany), assuming type II error probability of α=0.05, a power of 0.85, and an effect size of 0.730, based on previous data (pilot study). Adult (7-16months old) mixed-sex zebrafish (N=72; 40 for AB + 32 for TU) bred in the Animal Facility of i3S were used. They were kept in groups in 3.5 L tanks (maximum of 8 fish per liter), in a 14h:10h light:dark cycle, in a recirculating water system connected to a central unit of water purification and controlled conditions (27 ± 0.5 °C, pH of 7 + 0.5, conductivity of 800-900 μS). Adult fish were fed three times a day with a commercial diet (Zebrafeed 400-600 μm, Sparos, Olhão, Portugal). The same conditions of water were kept during the behavioural test.

### 2.3. Treatments

Zebrafish were randomly distributed to the control and amnesic treatment (positive control) group in each strain tested. Control group (N=37, 20 AB, 17 TU) was exposed to clean water, while amnesic treatment animals (N=35, 20 AB, 15 TU) were exposed to 20μM of MK-801 (MK-801 hydrogen maleate, M107, Sigma– Aldrich, USA) for 30 minutes.

### 2.4. Behavioral test - One trial inhibitory avoidance task

For this test, a modified passive avoidance task, based on the approach described by Blank et al. [7] to study memory and learning deficits was used. Our test is carried out in an apparatus that consists of a transparent tank surrounded by a 3D printed shell that is half white and half black with a moving barrier in the middle that can be raised or lowered manually allowing animals to visit both sides of the tank (Figure 1); this was connected to a shock device that modulates the shock from a power source (building instructions to be published). This paradigm consists on pairing an aversive experience with a previously neutral/preferred place [19]. The degree of avoidance is directly related to the latency of entering the previously preferred side and the time spent on each side after conditioning [20].

**Figure 1.**
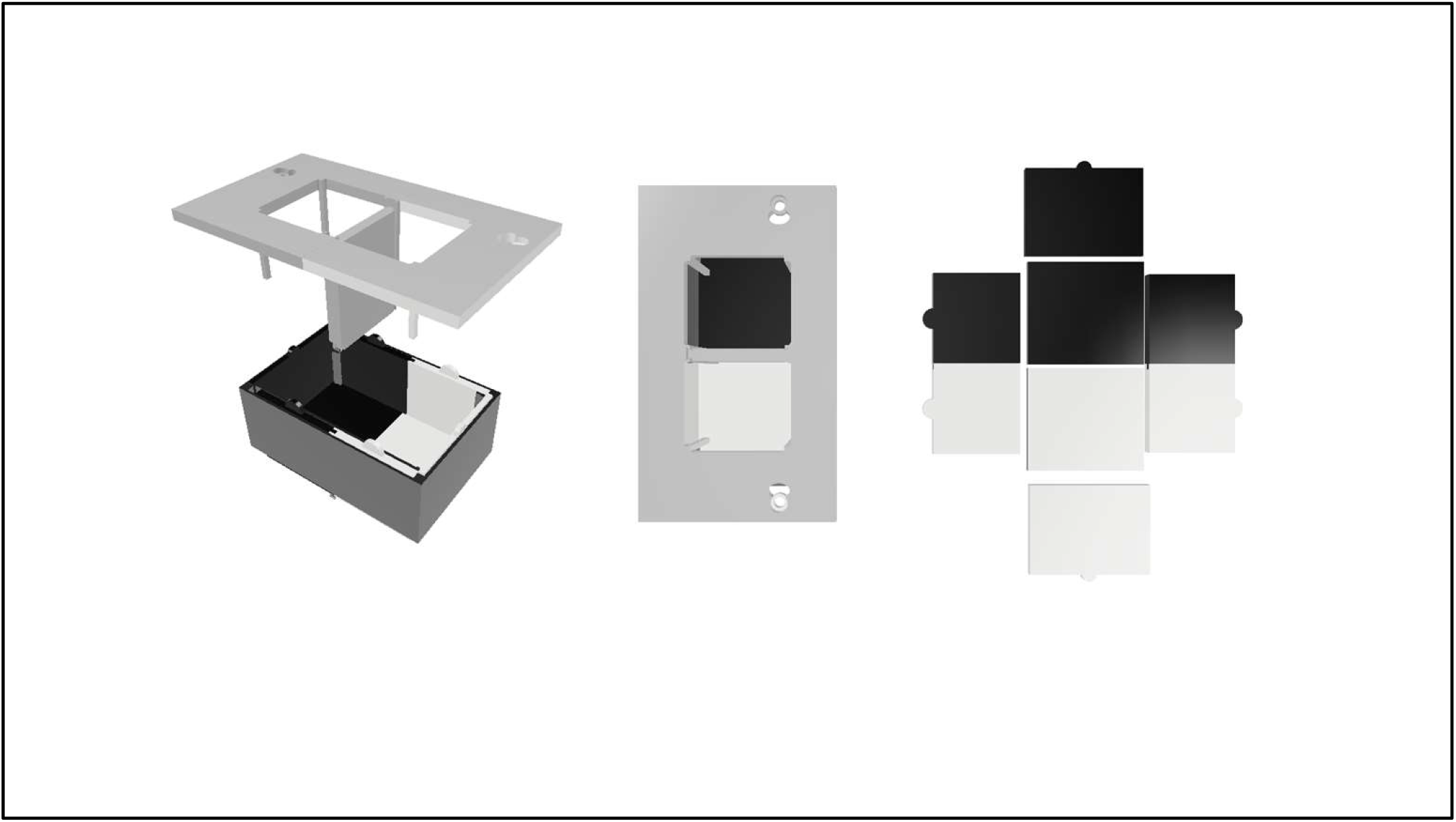
Schematic of the 3D printed apparatus in 3D rendering with an orthographic view (left), a top view (center) and a top view of the walls and floor separated from the apparatus (right). Holes and grooves on each side for electrode wiring and positioning and groove in the middle of the top part for the divider.

#### 2.4.1. Set-up

For the experiment, a custom-built 3D printed apparatus and custom-made hardware and software were made for the shock unit (Figure 1 and 2 and to be published). All test sessions were filmed from above with a camera (25 fps, Canon Legria HF R606, Ōta, Japan) that was fixed to a metal support at 45 cm of the apparatus, making sure no shadows were covering it.

**Figure 2.**
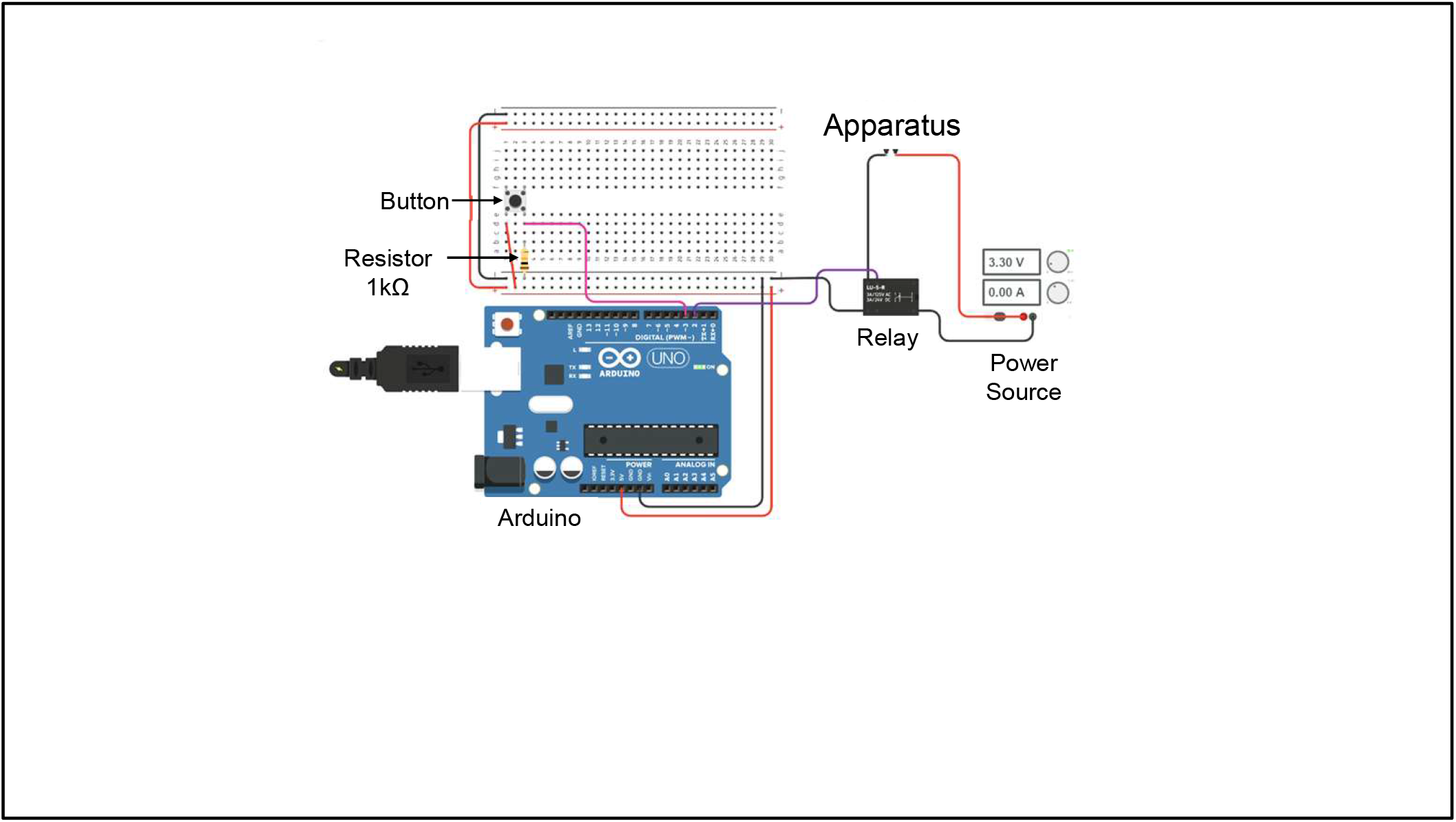
Schematic of the circuit layout of the shock unit

#### 2.4.2. Apparatus

This equipment consisted of an apparatus (Figure 1) with two equally sized black and white compartments. The apparatus has three parts: a Tecniplast TM plastic 1L breeding tank (16.5cm × 9.5cm), a shell that provides the desired colors to the apparatus (outer layer), and a shell that holds a divider and provides support for the electrodes (inner layer). The division of the compartments is done using a removable grey barrier with a thickness of 1 cm. The tank was filled with 300 ml of the recirculating water system (at 900 μS) where the zebrafish were housed, providing a 3 cm water column.

#### 2.4.3. Shock unit

This component consists of a direct current (DC) power supply (UNI-T, UTP1305, Uni-Trend Technology, Dongguan City, Guangdong), connected to relay controlled by an Arduino UNO ((Arduino, Ivrea, Italy); see annex for circuit layout and instructions will be published. We choose to use a DC constant current, instead of an alternating current because it is the most commonly used for electronic circuits while being the one allowing more control.

### 2.5. One-trial Inhibitory Avoidance test (Figure 3)

#### 2.5.1. Conditioning session

Individual fish were carefully placed on the white side where they remained for 1 min before the barrier was lifted. This was done to increase the ability of the animals to associate behavior-consequence during memory acquisition, in which temporal relations play an important role [21]. Thereafter the barrier was lifted 1 cm from the bottom of the tank, and immediately closed as soon as the animal crossed to the black side where an electric shock (3.3 ± 0.3V and 2A) was administered for 5 seconds after closing the barrier. Immediately after, the individual was then removed from the apparatus and placed in an amnesic bath of 20 μM of MK-801 (MK801 group) or in clean water (control group) for 30 min. The animals were then allowed to recover individually in clean water until the next day.

**Figure 3.**
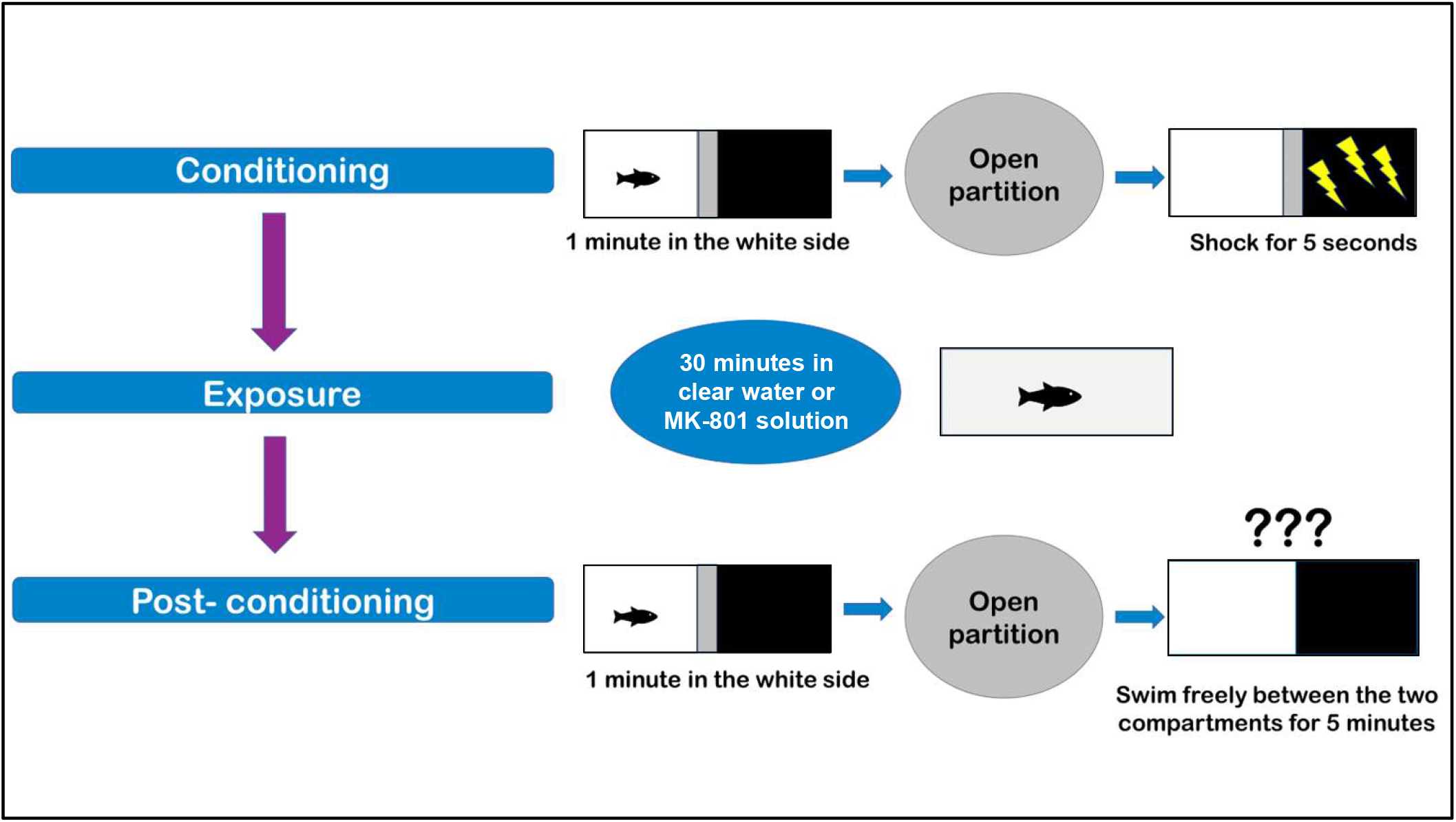
One-trial Inhibitory Avoidance Task behavior scheme. Control and MK-801 group were subjected to the same procedure, except they were individually placed in a tank with clear water or 20μM of MK-801 solution, respectively, for 30min.

#### 2.5.2. Postconditioning session

Twenty-four hours after the conditioning session, animals went back to the apparatus, and they were again retained for 1 min in the white zone. Subsequently, the barrier was lifted, and the animals were then able to swim for 5 minutes.

Latency to enter the black side, latency to reenter the white side, time spent on each side and frequency of crossings were measured.

Different electric shock characteristics (voltage and amperage) were preliminary tested until reaching the values presented here (to be published).

### 2.6. Video Analysis

All videos were analyzed by Ethowatcher, a free event-coding software developed by researchers from the Federal University of Santa Catarina, Brazil [22]. The analysis was carried out by an experienced researcher blind to the treatment, and two other blinded researchers confirmed the results by redoing the analysis.

### 2.7. Data Analysis

Data was analyzed using IBM SPSS™ 26 for Windows (SPSS Inc., Chicago, IL, USA) after exporting it to the Microsoft Excel™ 2010 (Microsoft Corporation, Redmond, WA, USA). First, data were analyzed to check their normality (Shapiro-Wilk test) and homoscedasticity, that is, homogeneity of variances between groups using the Levene test. If these assumptions were fulfilled, a parametric test was used; otherwise, the analysis was performed with a non-parametric test. For the parametric tests, a paired Student’s t-test (which compares means of continuous dependent variables between two related groups) was used for comparisons within groups, while the Wilcoxon signed-rank test is the correspondent nonparametric test. These tests were used to compare the differences within each group on the latency to enter the black side before and after the electric shock, and on the time spent between both sides of the apparatus during postconditioning.

For comparisons between groups, an independent Student’s t-test was used for parametric data, and a Mann-Whitney U test was used for nonparametric data. These tests compared the differences between groups regarding the latency to enter the black side before and after the electric shock, the time spent on each side of the apparatus, and the frequency of visits to the black side during postconditioning. All hypotheses were two-tailed tested and statistical significance was set at p ≤ 0.05.

## 3. Results

### 3.1. Animals

Two animals from the MK-801 group from the AB strain had to be eliminated from the analyses because there was a failure to close the barrier when individuals were introduced to the white side of the apparatus in the conditioning phase.

### 3.2. Treatments modulate the response of zebrafish after exposure

#### 3.2.1. Latency

First, in the conditioning phase, we checked if the animals behaved similarly before the treatment was applied; indeed, the latency to enter the black side were similar between control and MK801 group, and there were also no strain differences (p<0.05).

An animal that has learned and remembers the location of the aversive electric shock will present a higher latency to enter the black compartment after conditioning. Both control animals of the AB and TU strain took longer to enter the black side in the postconditioning compared with the conditioning phase. (Z=-2.315, p=0.021 and Z=2.062, p=0.039 for the AB and the TU strain, respectively). Both strains treated with the MK-801 did not show differences in this variable (Figure 4).

**Figure 4.**
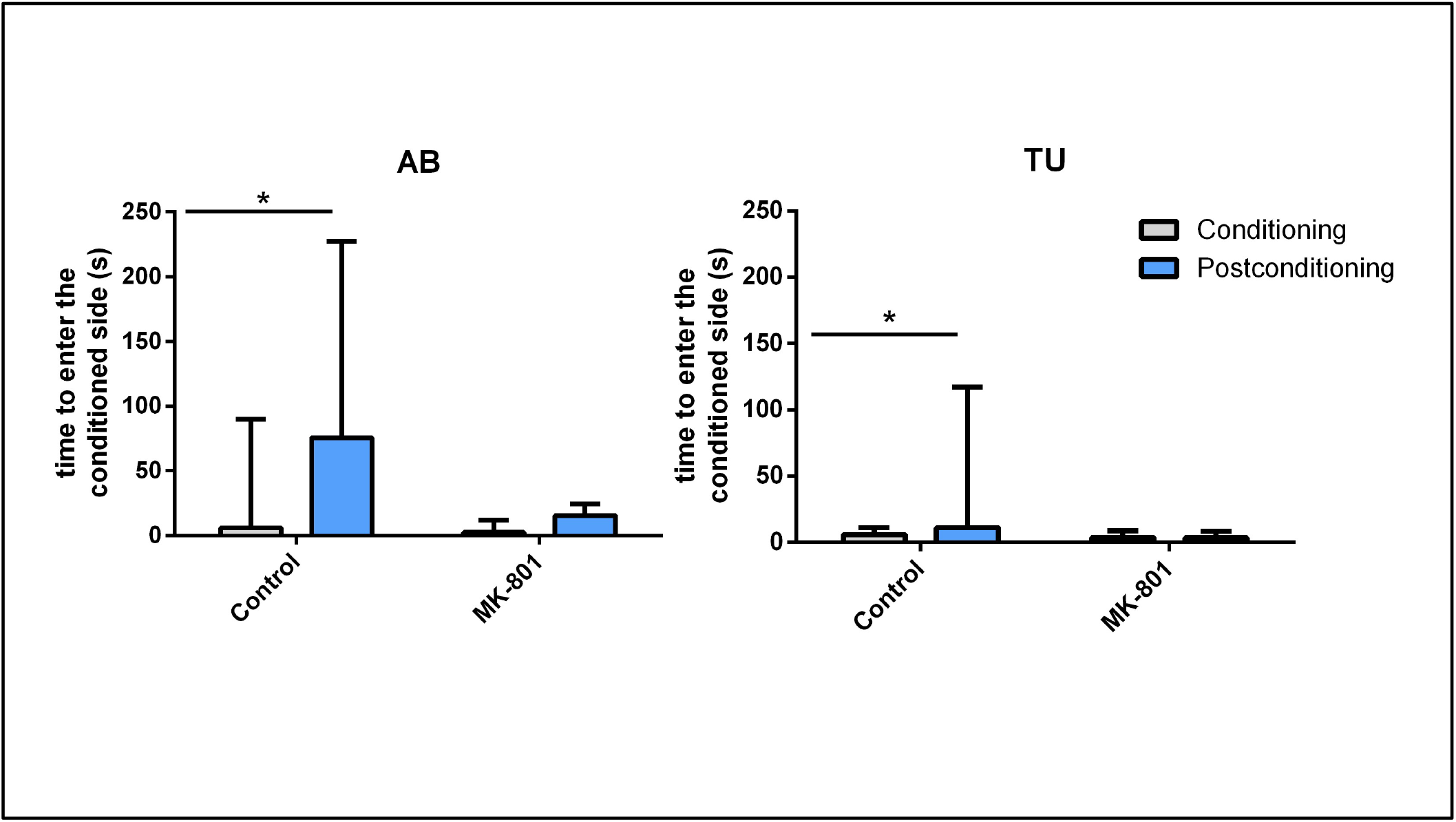
Latency to enter in the conditioned side in the conditioning phase and the post-conditioning phase for each treatment group and each strain. Control group: animals subjected to clean water (n= 20 for AB and 13 for TU); MK-801 group: animals subjected to 20 μM of MK-801 for 30 minutes (n= 20 for AB and 15 for TU). AB: animals from the WtAB strain, TU: animals from the WtTU strain. Data presented as median + IQR. * represents differences between phases within each group (p ≤0.039) using Wilcoxon signed-rank test.

#### 3.2.2. Time spent in each side

Regarding the time spent in each side of the apparatus during the postconditioning phase, the animals from the MK-801 group from both strains spent more time in the black side (Z=2.678, p=0.007 and Z=2.897, p=0.004, respectively). In the control group, animals from the TU strain control group spent more time in the black side (Z=2.271, p=0.023), whereas AB strain control showed no side preference (Figure 5).

**Figure 5.**
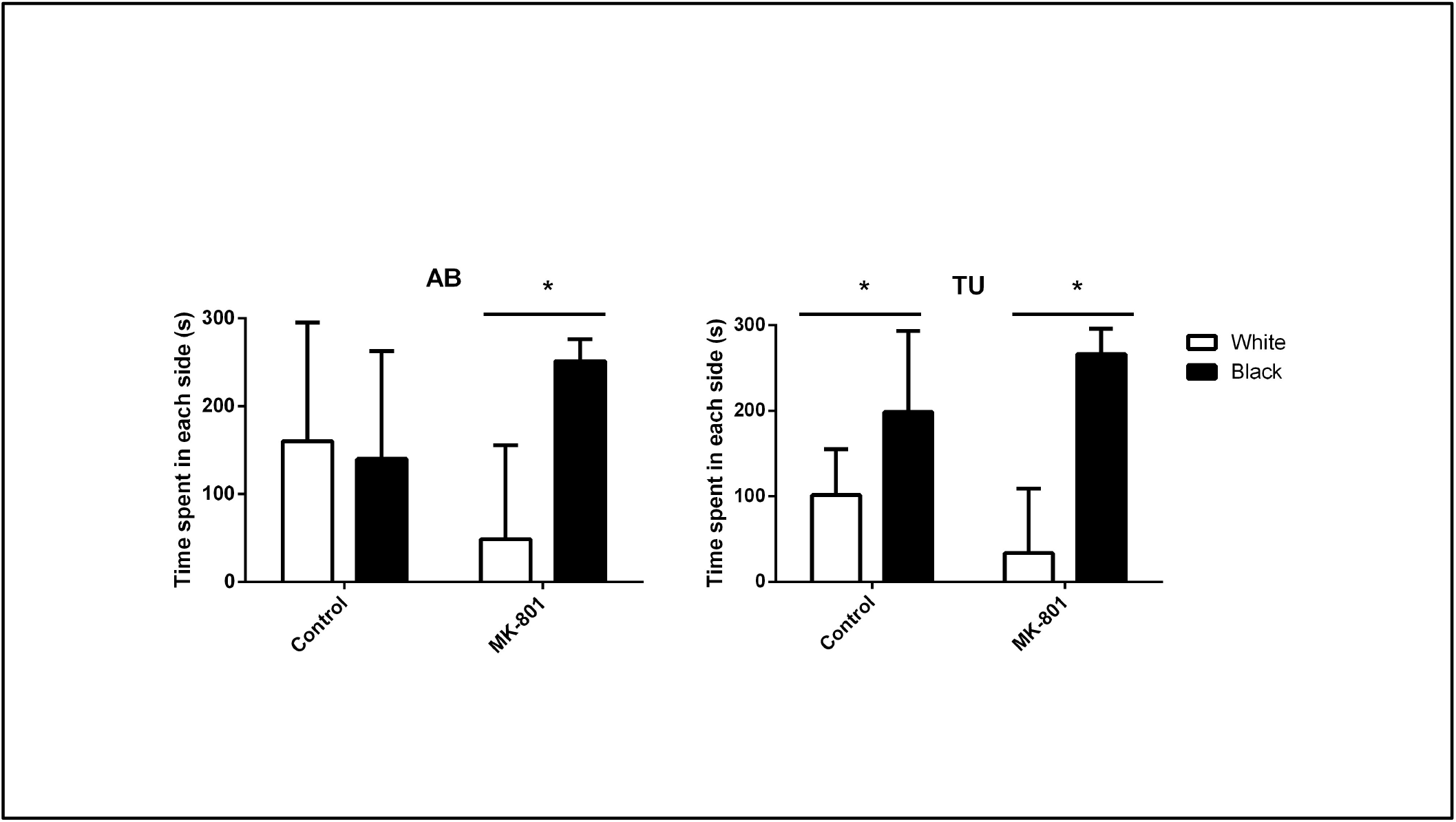
Time spent in each side of the apparatus in the post-conditioning phase for each treatment group and each strain. Control group: animals subjected to clean water (n= 20 for AB and 13 for TU); MK-801 group: animals subjected to 20 μM of MK-801 for 30 minutes (n= 20 for AB and 15 for TU). AB: animals from the WtAB strain, TU: animals from the WtTU strain. Data presented as median + IQR. * represents differences between sides of the apparatus within each group (p ≤0.023) using Wilcoxon signed-rank test.

#### 3.2.3. Frequency to visit each side

The frequency to visit the black side after the conditioning was reduced in the control group compared with the MK-801 group in the AB strain (p=0.017, U= 261.0); however, this difference was not detected in animals from the TU strain (Figure 6).

**Figure 6.**
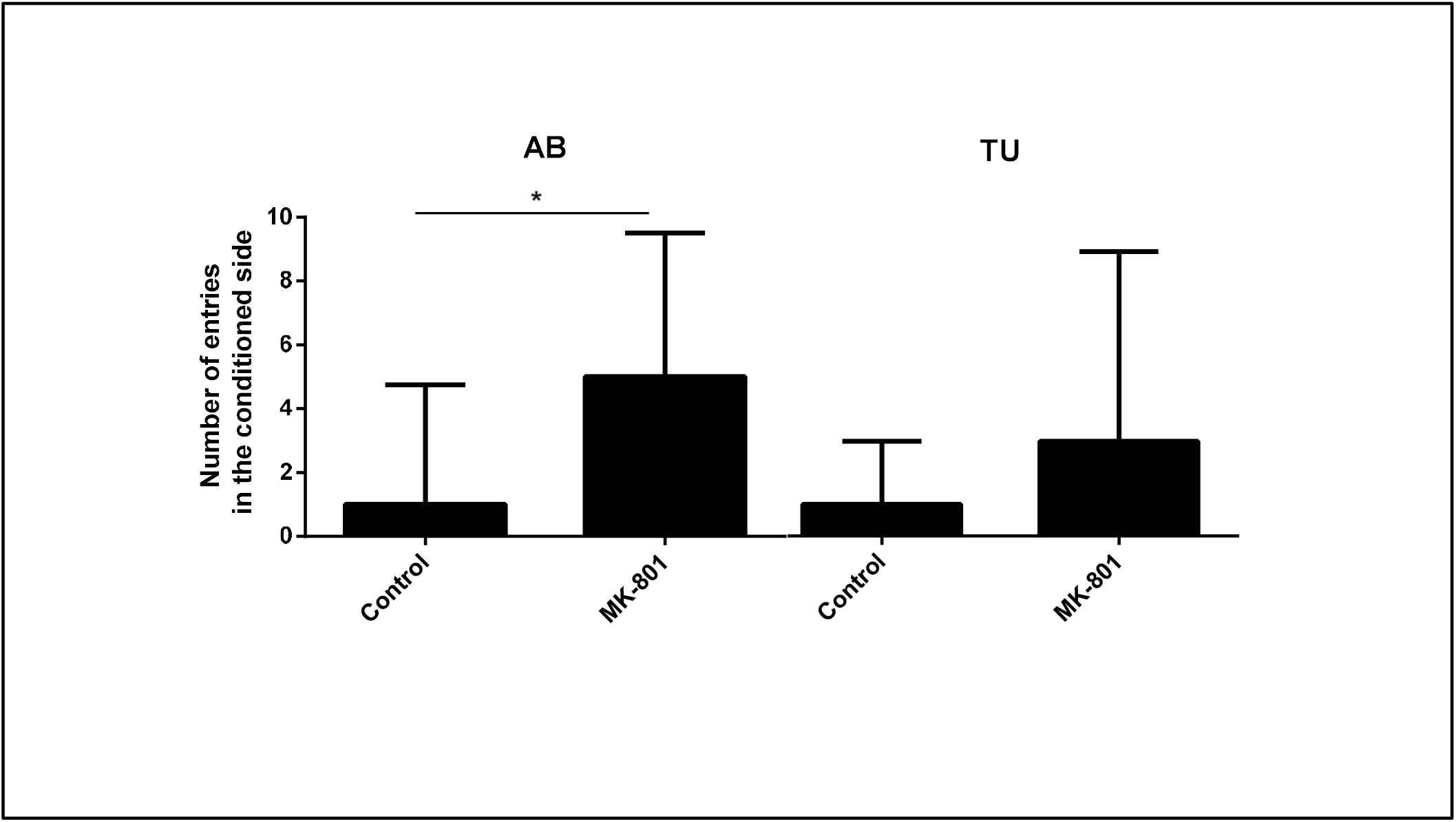
Number of entries in the conditioned side of the apparatus in the postconditioning phase for each treatment group and each strain. Control group: animals subjected to clean water (n= 20 for AB and 13 for TU) ; MK-801 group: animals subjected to 20 μM of MK-801 for 30 minutes (n= 20 for AB and 15 for TU). AB: animals from the WtAB strain, TU: animals from the WtTU strain. Data presented as median + IQR. * represents differences between groups in the AB strains (p<0.001) using Mann-Whitney test.

Although some results were different for the AB and TU strain regarding comparisons between the control and the MK-801 group in the postconditioning phase, no differences were detected between strains within these groups.

## 4. Discussion

When considering a new behavioural test or apparatus to use, replicability and reproducibility are of utmost importance as they are key points for our data to be compared with the ones from literature. In addition, accessibility and cost are two components that allow others to reproduce and replicate our data. This is important as then we can further create more accurate experiments and improve the standardization of the methodologies. Unfortunately, there is a methodology report crisis [2] mainly because the methods are not fully described, which leads to difficulties in replicating experiments. The test implemented in this work is an example of that, as the type of current and amperage was often not reported. To know the amperage to use is extremely essential, as the value determine the intensity of the shock. Additionally, there was also no fully available commercial version. Thus, we needed to build and test/ validate our own system. To validate this test, the electrical shock needed to create sufficient aversion for animals to avoid the conditioned side after just one trial, as in Blank’s work [7]. Additionally, the introduction of a drug known to interfere with memory would have to translate into data showing individuals that do not remember the shock, i. e., having the same latency to enter to the conditioned side before and after being subjected to the shock (conditioning). With both results being positive, we can say with confidence that our apparatus works.

In this study, using an apparatus fully assembled by us with refined parameters to the electrical shock, animals from both strains behaved similarly to the ones used by Blank and colleagues [7]; the control animals took longer to enter the black side after conditioning. Our MK-801 animals took the same time to go to the black side before and after the shock, meaning that the animals did not remember the shock, therefore not displaying learned aversion to the black side. These results already show two important premises: the shock had induced enough aversion in the control animals of both strains, and the NMDA antagonist disrupted the memory of the electrical shock in both strains.

Different zebrafish strains may have different behavioural profiles, which may result on different results of the test we want to validate [23]. However, we found no differences between the TU and AB strains tested in each measured variable.

Nevertheless, the behavioural profile of each strain was different with respect to the time spent in each side of the tank and the frequency of entries in the black side. During postconditioning, the AB control did not reveal a preference for any side of the tank, while AB MK-801 group spent more time in the black side. These results are in line with the ones from the latency to enter the black side, where control animals showed avoidance for the black side, but not the MK-801 animals (impaired memory). On the contrary, the TU control animals spent more time in the previously preferred place, the black side. It has been described that TU strain recovered faster its normal activity in the novel tank test, indicating less anxiety-like behaviour in space occupation than the AB strain [16], which could explain the results obtained. Thus, the TU control may have low levels of anxiety compared with AB control, allowing them to recover from the previously unpleasant experience and spending more time in the black side (the preferred side) where they no longer receive an electric shock. Similarly, AB control entered less in the black side than the AB MK801 group, while the MK-801 and control group from the TU presented equal number of entries in this side. These results did not indicate that the TU strain did not learn, as they remembered the shock, showing longer time to enter the black side after conditioning, as referred. Also, the amnesic group MK-801 was only different from the control group in both strains in the latency to enter the black side. Thus, the latency to enter the black side before and after conditioning appears to be the most important variable to evaluate memory in this task, while the others may be influenced by anxiety.

Except for the frequency to enter the black side, both strains had the same response to the amnesiac drug, showing that the NMDA receptors were similarly affected by MK-801, and both strains forgot the aversive experience.

Avoidance tasks are very important in neuroscience as they have been used for a variety of assessments, from learning to anxiety [12, 24, 25]. This is based on the assessment of a nonspecific distress response, the escape response, which is a key behavior for the survival of species, and thus a well conserved trait ([26]). In this test apparatus, initially, the zebrafish have the tendency to avoid the white side, preferring to go to the black compartment, while the white color may expose the animals to potential threats; different anxiety levels may modulate this behavior. Later (postconditioning) the animal must passively avoid a real aversive stimulus, not going to the previously preferred place, fear is potentially triggered but fades during the phase and the animal enters the black side. In this sense, anxiety may also be involved in the animals’ decision [27].

Linked with this, the animal needs to remember previous unpleasant experiences to perform the task and avoid another potential electric shock; for that, learning and memory processes are key. Since memory is mediated by synaptic plasticity [28, 29], we can look at neurotransmitter receptors and modulate them with amnesic drugs such as*N*-methyl-D-aspartate (NMDA) receptor has been reported to be involved in fear-avoidance learning [27]. Previous studies in zebrafish and other species [7, 30-32] have shown that MK-801 is effective in preventing memory formation when given after training in similar inhibitory avoidance tasks [7, 32].

Fortunately, developments in 3D printing technology made the building of this type of tests very affordable and the tools are easy to access when paired with the availability of open-source hardware microcontrollers. This inexpensive easy to assemble apparatus can be used to perform a reproducible task for memory and learning assessment in adult zebrafish from both AB and TU strains, as we provide here all the details to build the test, and a pharmacological validation.

## Author Contributions

Conceptualization: A.M.V., J.F.; Methodology: A.M.V., J.F., P.F.; Validation: A.M.V., J.F., S.B., J.S; Investigation: J.F., S.B., J.S., I.C.; Formal analysis: A.M.V., J.F.; Resources: A.M.V. and I.A.S.O.; Writing—original draft preparation: A.M.V. and J.F.; Writing—review and editing: A.M.V., I.A.S.O., I.C., P.F.; Supervision: A.M.V.; Project administration: A.M.V.; Funding acquisition: A.M.V. and I.A.S.O. All authors have read and agreed to the published version of the manuscript.

## References

[1] E.E. Commission, report on the statistics on the use of animals for scientific purposes in the Member States of the European Union in 2015-2017, Brussels 5 (2020).

[2] C.M. Loss, F.F. Melleu, K. Domingues, C. Lino-de-Oliveira, G.G. Viola, Combining Animal Welfare With Experimental Rigor to Improve Reproducibility in Behavioral Neuroscience, Front Behav Neurosci 15 (2021).

[3] W.M.S. Russell, R.L. Burch, The principles of humane experimental technique, Methuen 1959.

[4] J.M. Ferreira, I.A.S. Olsson, A.M. Valentim, Disentangling the terminology of the light/dark and white/black test in zebrafish, (2023).

[5] J.D. Sweatt, Rodent behavioral learning and memory models, Mechanisms of Memory, 2nd ed. Elsevier, UK (2010) 77–103.

[6] B. Jänicke, H. Coper, Tests in rodents for assessing sensorimotor performance during aging, Advances in Psychology, Elsevier 1996, pp. 201–233.

[7] M. Blank, L.D. Guerim, R.F. Cordeiro, M.R. Vianna, A one-trial inhibitory avoidance task to zebrafish: rapid acquisition of an NMDA-dependent long-term memory, Neurobiology of learning and memory 92(4) (2009) 529–534.

[8] A. Facciol, M. Iqbal, A. Eada, S. Tran, R. Gerlai, The light-dark task in zebrafish confuses two distinct factors: Interaction between background shade and illumination level preference, Pharmacology Biochemistry and Behavior 179 (2019) 9–21.

[9] S. British Association for the Advancement of, Report of the British Association for the Advancement of Science, London, 1831.

[10] P. Yang, R. Kajiwara, A. Tonoki, M. Itoh, Successive and discrete spaced conditioning in active avoidance learning in young and aged zebrafish, Neuroscience Research 130 (2018) 1–7.

[11] L. Truong, D. Mandrell, R. Mandrell, M. Simonich, R.L. Tanguay, A rapid throughput approach identifies cognitive deficits in adult zebrafish from developmental exposure to polybrominated flame retardants, Neurotoxicology 43 (2014) 134–142.

[12] X. Xu, T. Scott-Scheiern, L. Kempker, K. Simons, Active avoidance conditioning in zebrafish (Danio rerio), Neurobiology of learning and memory 87(1) (2007) 72–77.

[13] R. Manuel, M. Gorissen, C. Piza Roca, J. Zethof, H.v.d. Vis, G. Flik, R.v.d. Bos, Inhibitory avoidance learning in zebrafish (Danio rerio): effects of shock intensity and unraveling differences in task performance, Zebrafish 11(4) (2014) 341–352.

[14] R. Aoki, T. Tsuboi, H. Okamoto, Y-maze avoidance: An automated and rapid associative learning paradigm in zebrafish, Neuroscience research 91 (2015) 69–72.

[15] M. Gorissen, R. Manuel, T.N. Pelgrim, W. Mes, M.J. de Wolf, J. Zethof, G. Flik, R. van den Bos, Differences in inhibitory avoidance, cortisol and brain gene expression in TL and AB zebrafish, Genes, Brain and Behavior 14(5) (2015) 428–438.

[16] C. Vignet, M.-L. Bégout, S. Péan, L. Lyphout, D. Leguay, X. Cousin, Systematic screening of behavioral responses in two zebrafish strains, Zebrafish 10(3) (2013) 365–375.

[17] M. Sison, R. Gerlai, Associative learning performance is impaired in zebrafish (Danio rerio) by the NMDA-R antagonist MK-801, Neurobiology of learning and memory 96(2) (2011) 230–237.

[18] M. Sison, R. Gerlai, Behavioral performance altering effects of MK-801 in zebrafish (Danio rerio), Behavioural brain research 220(2) (2011) 331–337.

[19] A.M. Stewart, O. Braubach, J. Spitsbergen, R. Gerlai, A.V. Kalueff, Zebrafish models for translational neuroscience research: from tank to bedside, Trends in neurosciences 37(5) (2014) 264–278.

[20] G.D. Readman, S.F. Owen, J.C. Murrell, T.G. Knowles, Do fish perceive anaesthetics as aversive?, PLoS One 8(9) (2013) e73773.

[21] T. Abel, K.M. Lattal, Molecular mechanisms of memory acquisition, consolidation and retrieval, Current opinion in neurobiology 11(2) (2001) 180–187.

[22] C.F.C. Junior, C.N. Pederiva, R.C. Bose, V.A. Garcia, C. Lino-de-Oliveira, J. Marino-Neto, ETHOWATCHER: validation of a tool for behavioral and video-tracking analysis in laboratory animals, Computers in biology and medicine 42(2) (2012) 257–264.

[23] G. Audira, P. Siregar, S.-A. Strungaru, J.-C. Huang, C.-D. Hsiao, Which zebrafish strains are more suitable to perform behavioral studies? A comprehensive comparison by phenomic approach, Biology 9(8) (2020) 200.

[24] R.M. Colwill, M.P. Raymond, L. Ferreira, H. Escudero, Visual discrimination learning in zebrafish (Danio rerio), Behavioural Processes 70(1) (2005) 19–31.

[25] F.E. Williams, D. White, W.S. Messer Jr, A simple spatial alternation task for assessing memory function in zebrafish, Behavioural Processes 58(3) (2002) 125–132.

[26] D.A. Evans, A.V. Stempel, R. Vale, T. Branco, Cognitive control of escape behaviour, Trends in cognitive sciences 23(4) (2019) 334–348.

[27] M. La-Vu, B.C. Tobias, P.J. Schuette, A. Adhikari, To approach or avoid: an introductory overview of the study of anxiety using rodent assays, Front Behav Neurosci 14 (2020) 145.

[28] G. Neves, S.F. Cooke, T.V. Bliss, Synaptic plasticity, memory and the hippocampus: a neural network approach to causality, Nature Reviews Neuroscience 9(1) (2008) 65–75.

[29] S.J. Martin, P.D. Grimwood, R.G. Morris, Synaptic plasticity and memory: an evaluation of the hypothesis, Annual review of neuroscience 23(1) (2000) 649–711.

[30] R. Burchuladze, S.P. Rose, Memory formation in day-old chicks requires NMDA but not non-NMDA glutamate receptors, European Journal of Neuroscience 4(6) (1992) 533–538.

[31] K. Liang, M.-H. Lin, Y.-M. Tyan, Involvement of amygdala N-methyl-D-asparate receptors in long-term retention of an inhibitory avoidance response in rats, The Chinese journal of physiology 36(1) (1993) 47–56.

[32] C. Castellano, V. Cestari, A. Ciamei, F. Pavone, MK-801-induced disruptions of one-trial inhibitory avoidance are potentiated by stress and reversed by naltrexone, Neurobiology of Learning and Memory 72(3) (1999) 215–229.

